# Fix your membrane receptor imaging: Actin cytoskeleton and CD4 membrane organization disruption by chemical fixation

**DOI:** 10.1101/450635

**Authors:** Pereira Pedro M., David Albrecht, Caron Jacobs, Mark Marsh, Jason Mercer, Ricardo Henriques

**Affiliations:** MRC-Laboratory for Molecular Cell Biology. University College London, London, UK; Department of Cell and Developmental Biology, University College London, London, UK; The Francis Crick Institute, London, UK; Current address: Gene Expression & Biophysics Group, Division of Chemical Systems & Synthetic Biology, Institute for Infectious Disease & Molecular Medicine (IDM), Department of Integrative Biomedical Sciences, Faculty of Health Sciences, University of Cape Town

**Author notes:** Equal contributing authors (Pereira Pedro M., David Albrecht).

**Keywords:** Super-resolution microscopy, Single Molecule Localization Microscopy, NanoJ-Fluidics, NanoJ-SQUIRREL, Clustering analysis, Fixation, Artefacts, Actin cytoskeleton, CD4 membrane receptor

## Abstract

Single-molecule localization microscopy (SMLM) techniques allow near molecular scale resolution (~ 20nm) as well as precise and robust analysis of protein organization at different scales. SMLM hardware, analytics and probes have been the focus of a variety of studies and are now commonly used in laboratories across the world. Protocol reliability and artefact identification are increasingly seen as important aspects of super-resolution microscopy. The reliability of these approaches thus requires in-depth evaluation so that biological findings are based on solid foundations. Here we explore how different fixation approaches that disrupt or preserve the actin cytoskeleton affect membrane protein organization. Using CD4 as a model, we show that fixation-mediated disruption of the actin cytoskeleton correlates with changes in CD4 membrane organization. We highlight how these artefacts are easy to overlook and how careful sample preparation is essential for extracting meaningful results from super-resolution microscopy.

## Introduction

Super-resolution microscopy is a fundamental tool for exploring and understanding nanoscale biological assemblies. Single-molecule localization microscopy (SMLM) techniques in particular, such as photoactivated localization microscopy (PALM) (1) and stochastic optical reconstruction microscopy (STORM) (2), are the optical imaging gold standards to study membrane protein organization (3). SMLM techniques provide high spatial resolution (~20 nm) and allow for statistical, nonbiased analysis of membrane protein nanoscale organizations (1, 2, 4, 5). Thereby, super-resolution microscopy has provided new views on the organization of membrane receptors, from immune sensing to pathogen engagement (6). The organization of receptors into micro- and nanoclusters at the plasma membrane is a common feature and an important regulatory mechanism for cell signaling and activation (7–11). Thus, analyzing the nanoscale level organization of these molecules is critical to understand basic regulation of cellular signaling but also to understand the function of these proteins in disease. For example, CD4 plays an important role in immune cell activation through its ability to enhance T-cell receptor (TCR)-mediated signaling by binding to the antigen-presenting major histocompatibility complex II (MHCII) (12). Besides its importance in immune signaling, CD4 is also the primary cellular receptor for human immunodeficiency viruses (HIV) (12, 13). The importance of super-resolution in the study of membrane receptor organization and function cannot be overstated. A recent example is the characterization of the spatiotemporal dynamics and stoichiometry of the interactions between CD4 (and co-receptors) and HIV-1 in the context of viral entry, impossible to achieve without molecular imaging approaches (13).

A key component of membrane organization is the actin cytoskeleton (14, 15). The actin cortex underlies the plasma membrane and interacts both with lipids and membrane proteins, functioning as a dynamic scaffold providing support and force for the continuous remodelling of membrane receptor organization (16–18). It is not surprising that the actin cytoskeleton has been the subject of a considerable number of studies in a variety of biological settings, from viral engagement to axon organization using (super-resolution) microscopy (16, 19, 20).

The increased resolution and detailed analytic information provided by SMLM requires rigorous scrutiny of collected data (21–23). The succession of steps from the native organization of a receptor in the plasma membrane to the final super-resolution image can be significantly influenced by artefacts, particularly if imaging requires chemical fixation (21–23). Ideally, chemical fixation preserves the macroscopic structure of the sample as well as the native nanoscale organization of target proteins. However, true preservation at the subcellular level is not trivial, as known from electron microscopy studies (24). Furthermore, chemical fixation does not immediately immobilize membrane-associated proteins (25). Thus, given the increase in resolution afforded by super-resolution microscopy, the effect of fixation has been the focus of several recent studies (21–23). Importantly, there are multiple chemical fixation methods, differing by the fixative used (e.g. paraformaldehyde, glutaraldehyde, glyoxal or methanol), the buffer composition (e.g. phosphate buffered saline, cytoskeleton stabilizing buffer or PIPES-EGTA-magnesium buffer), and physical conditions (temperature and duration) (21–23, 26, 27). There is, at this stage, no standardized sample preparation protocol to study membrane protein organization. Moreover, to the best of our knowledge, there is no correlative study to understand how, in the same cells, fixation-induced changes in the actin cytoskeleton may affect membrane protein organization.

Here, we analyze how the morphology of the actin cytoskeleton changes with different chemical fixation protocols and how these changes correlate with the membrane organization of the membrane receptor CD4 (Fig. 1). Conditions that have detrimental effects on cytoskeleton organization correlate with changes in the membrane organization of CD4. We suggest that careful sample preparation and handling during all steps leading to the final image is essential for all scientists.

**Fig. 1.**
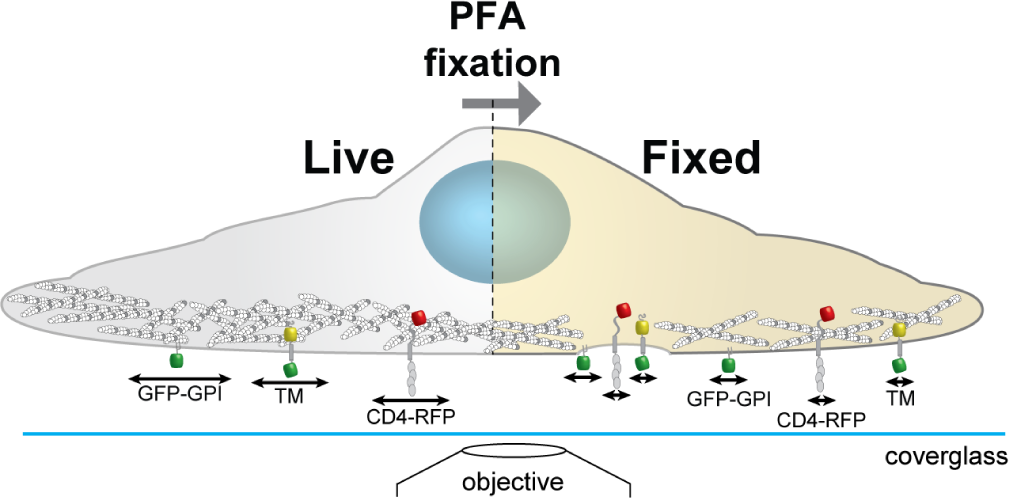
Schematics of the experimental workflow to correlate actin morphology with CD4 membrane organization. We analyse on the same cells how the actin cytoskeleton morphology changes with different chemical fixation protocols and how this correlates with the membrane organization and mobility of CD4. Cortical actin (white and grey circles); arrows represent protein mobility; GPI anchored GFP (GFP-GPI); artificial transmembrane protein with cytosolic and extracellular domains (mHoneydew and YFP, respectively - TM); CD4 fused to TagRFP-T (CD4-RFP).

## Suboptimal fixation protocols affect the actin cytoskeleton and CD4 membrane organization differently

To understand the effect of suboptimal actin fixation protocols on CD4 membrane organization we correlated live-cell and fixed-cell actin and CD4 organization using NanoJ-Fluidics (28) (Fig. 2a) and Structured Illumination Microscopy (SIM) (29). We imaged actin in live COS7 cells with an utrophin domain (UtrCH-GFP) (30) probe and CD4 tagged with TagRFP-T. We performed chemical fixation using three different chemical fixation protocols, 4% paraformaldehyde (PFA) in PBS at 23°C, 4% PFA in PEM(22) at 4°C or at 37°C (Fig. 2b-d). Subsequently, using NanoJ-SQUIRREL (21), to compare the live-cell vs fixed-cell organization of actin and CD4, we were able to identify the effects of the suboptimal (4% PFA in PBS at 23°C and 4% PFA in PEM at 4°C) fixation protocols on these targets and compare with the optimal protocol (37°C 4% PFA in PEM) (22). As expected (22, 23), using PBS we observed a striking disruption of the actin cytoskeleton (Fig. 2b). The fixation resulted in an almost indiscernible actin cytoskeleton, which translates to a NanoJ-SQUIRREL error map exhibiting strong artefacts (Fig. 2b). Using PEM buffer, more suited for actin preservation (22), but at a suboptimal temperature (4°C), we see a much milder disruption of the actin organization (Fig. 2c). Pre-warming the PFA-containing PEM buffer to 37°C yielded a similar difference between live and fixed sample as measured by NanoJ-SQUIRREL (Fig. 2d). Regardless of the fixation approach we did not see an effect on CD4 membrane organization, quantified on the error maps where most of the differences are due to vesicle motion during fixation (Fig. 2b-d).

**Fig. 2.**
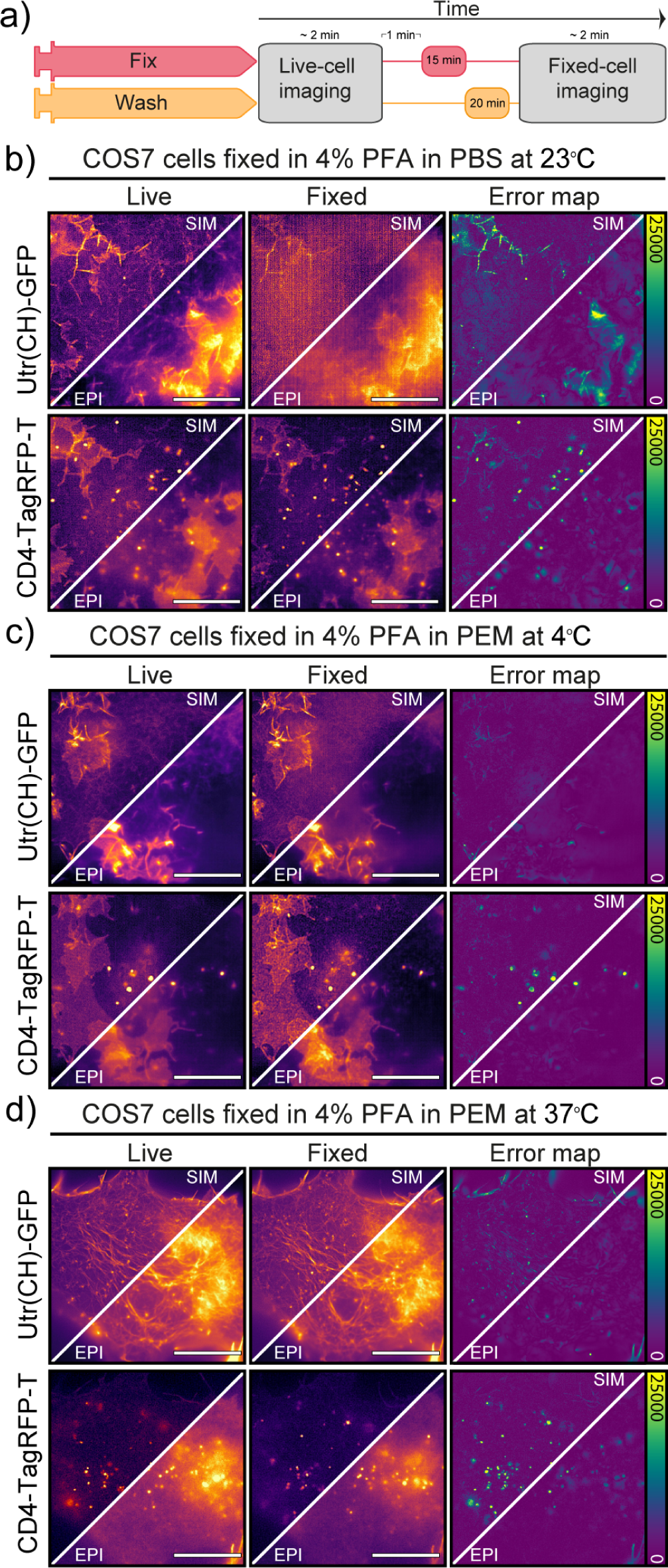
Effect of suboptimal fixation conditions on actin and CD4 organization. a) NanoJ-Fluidics protocol to perform the live-to-fixed cell correlation under different suboptimal fixation conditions. b) Epifluorescence and SIM imaging of COS7 cell expressing Utr(CH)-GFP and CD4-TagRFP-T live (Live) and fixed (Fixed) with 4% PFA in PBS at 23°C and corresponding NanoJ-SQUIRREL error maps (Error map). c) same as in b) but fixation was performed with 4% PFA in PEM at 4°C. d) same as in b) but fixation was performed with 4% PFA in PEM at 37°C. Scale bars are 10 µm.

## The fixation protocol influences CD4 cluster size and cluster density at the cell surface

To ascertain if CD4 membrane organization was correlated with fixation-mediated actin cytoskeleton disruption we repeated the live-to-fixed cell correlation using SMLM and PEM with different fixation temperatures. PEM is an ideal buffer for actin preservation (22), and the range of temperatures provide a different fixation efficiencies, with decreasing efficiency from 37°C (ideal) to 23°C (intermediate) to 4°C (lowest efficiency). We took advantage of the versatile NanoJ-Fluidics (28) framework to correlate live and fixed cell imaging of COS7 cells (Fig. 3a). As expected, regardless of the fixation strategy we obtain a fairly homogeneous distribution of CD4 on the surface of COS7 cells (Fig. 3b) at an in-cell high-resolution (43-50 nm by FRC (21)). To further explore the nature of the CD4 organization we used SR-Tesseler (4) to determine if the cluster sizes and cluster density of CD4 would change depending on the fixation approach (Fig. 3c). Interestingly, despite the little changes observed by SIM (Fig. 2), both CD4 cluster size and cluster density changed with the fixation approach. Whereas the mean CD4 cluster size in ideal conditions (PEM buffer at 37°C) is 59 nm, reducing the temperature to 23°C or 4°C is enough to change CD4 organization, increasing the mean cluster size to 65 nm (p<0.001). The fixation conditions also influence the CD4 cluster density in COS7 cells, with densities of 1.3 clusters/µm^2^ for 37°C, 1.8 clusters/µm^2^for 23°C and 3.8 clusters/µm^2^ for 4°C.

**Fig. 3.**
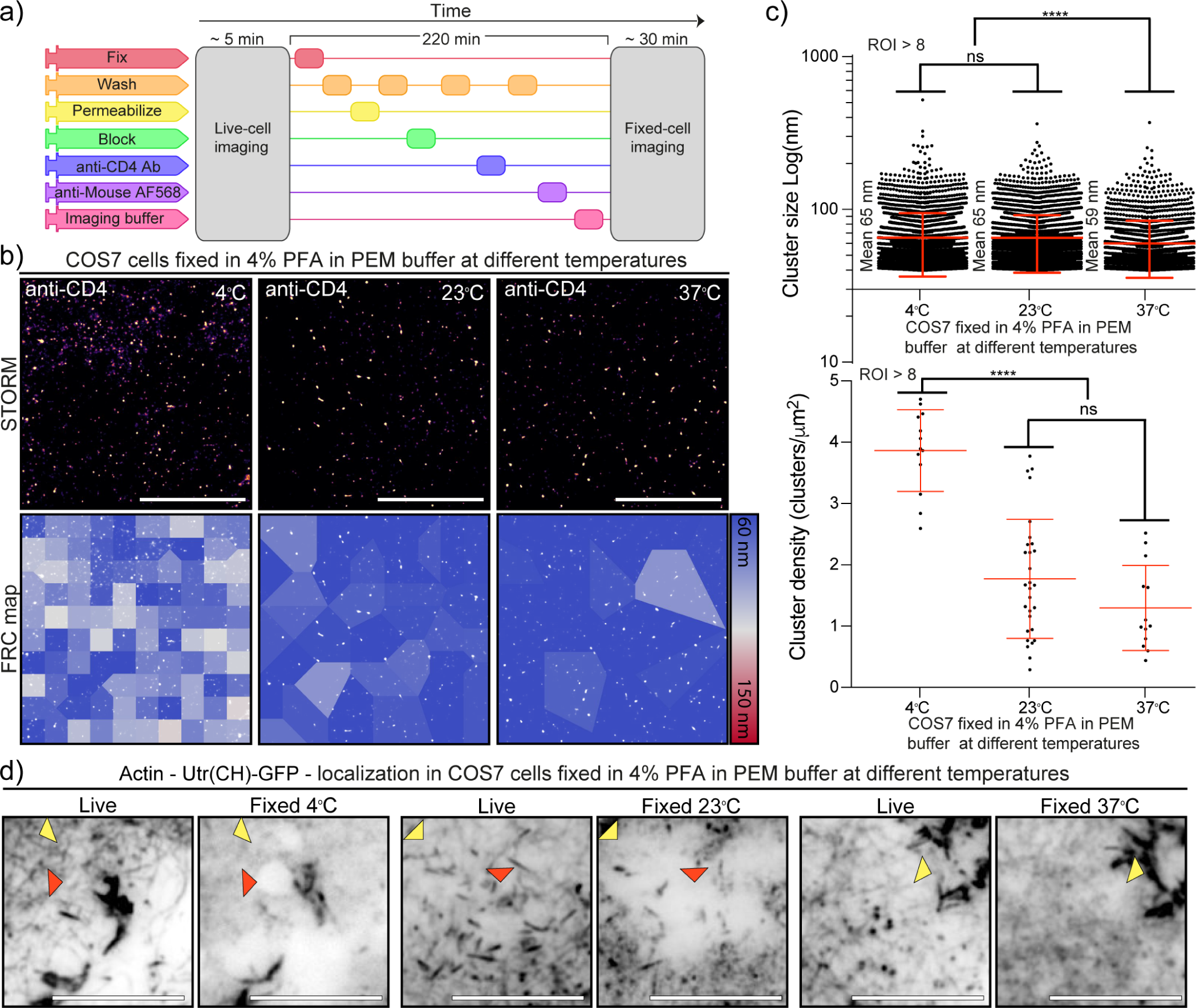
Correlation between CD4 membrane organization and actin structure fidelity upon fixation. a) NanoJ-Fluidics protocol to perform the live-to-fixed cell correlation under different fixation conditions. b) CD4 STORM imaging after fixation in different conditions (Top) and FRC map of the same region (Bottom). Scale bars are 5 µm. c) CD4 cluster size and cluster density under different fixation conditions. d) Diffraction limited (TIRF) live-to-fixed cell correlation using different fixation conditions. Red arrowheads highlight areas where actin disappeared upon fixation. Yellow arrowheads highlight areas where there is a difference in actin organization due to fixation. Scale bars are 1 µm.

## Fixation-induced CD4 organization changes correlate with actin cytoskeleton preservation

We posited that fixation-induced changes in CD4 organization could be related to disruption of the actin cytoskeleton (Fig. 2). To determine if the actin cytoskeleton was affected we compared the actin organization in the cells pre- and post-fixation (Fig. 3d). We observed a disruption of the actin cytoskeleton at 23°C and 4°C when compared with fixation at 37°C (Fig. 3d). Independent of the fixation condition the post-fixation actin organization is different from the live-cell actin organization (Fig. 3d yellow arrowheads). With the decrease in fixation temperature there is a step-wise decrease in the fidelity of the fixed-cell actin structure in relation to the one observed in live-cells. At lower fixation temperatures, actin filaments disappear and there are gaps in the actin structure, possibly related to cell detachment from the substrate or actin cytoskeleton disruption (Fig. 3d red arrowheads). These artefacts are less prevalent in cells fixed under conditions that preserve the actin cytoskeleton structure.

## The changes in CD4 membrane organization are not related to fixation-induced cell membrane disruption as seen by single-particle tracking

The difference in membrane receptor organization could be the result of fixation efficiency dependence on temperature. Employing our live-to-fix approach, we sought to determine how quickly the addition of PFA-containing PEM buffer immobilizes membrane associated proteins (Fig. 4). An artificial trans-membrane protein with a~ 30 kDa cytosolic and extracellular domain (mHoneydew and YFP, respectively) was expressed in COS7 cells and individual proteins tracked with uPAINT (31), i.e. by adding low concentration (~20 pMol) of Atto647N labeled anti-GFP nanobodies (Chromotek) to the medium (Fig. 4b, first panel). Diffusion coefficients based on particle velocity were 0.27 ±0.06 µm^2^/s (mean). Exchanging the cell culture medium with 37°C pre-warmed 4% PFA in PEM immediately reduced the diffusion speed of transmembrane proteins (Fig. 4b, middle panel, arrow) and, after 10 minutes fixation, 97% of proteins were immobilized (D < 0.05 µm^2^/s) (Fig. 4c). Addition of cold (4°C) 4% PFA in PEM had similar effects on measured diffusion coefficients and mobility (Fig. 4c). Next, we tested if the same was true for a GPI-anchored protein that lacks any cytosolic domain that might interact directly with the cytoskeleton. GPI-anchored GFP was tracked via anti-GFP nanobodies. Addition of warm (37°C) or cold (4°C) 4% PFA in PEM buffer to live cells reduced the mobility of tracked individual particles without immobilizing them completely (Fig. 4d, middle panel, arrow). In contrast to the transmembrane probe, only some particles were immobilized after fixation for 10 minutes. The reduction of diffusion coefficients as measured by velocity or mean square displacement (Fig. 4e, left and middle panel) was not significantly different based on temperature. The mobile fraction was reduced to 36% and 32% (mean) after warm and cold fixation, respectively.

**Fig. 4.**
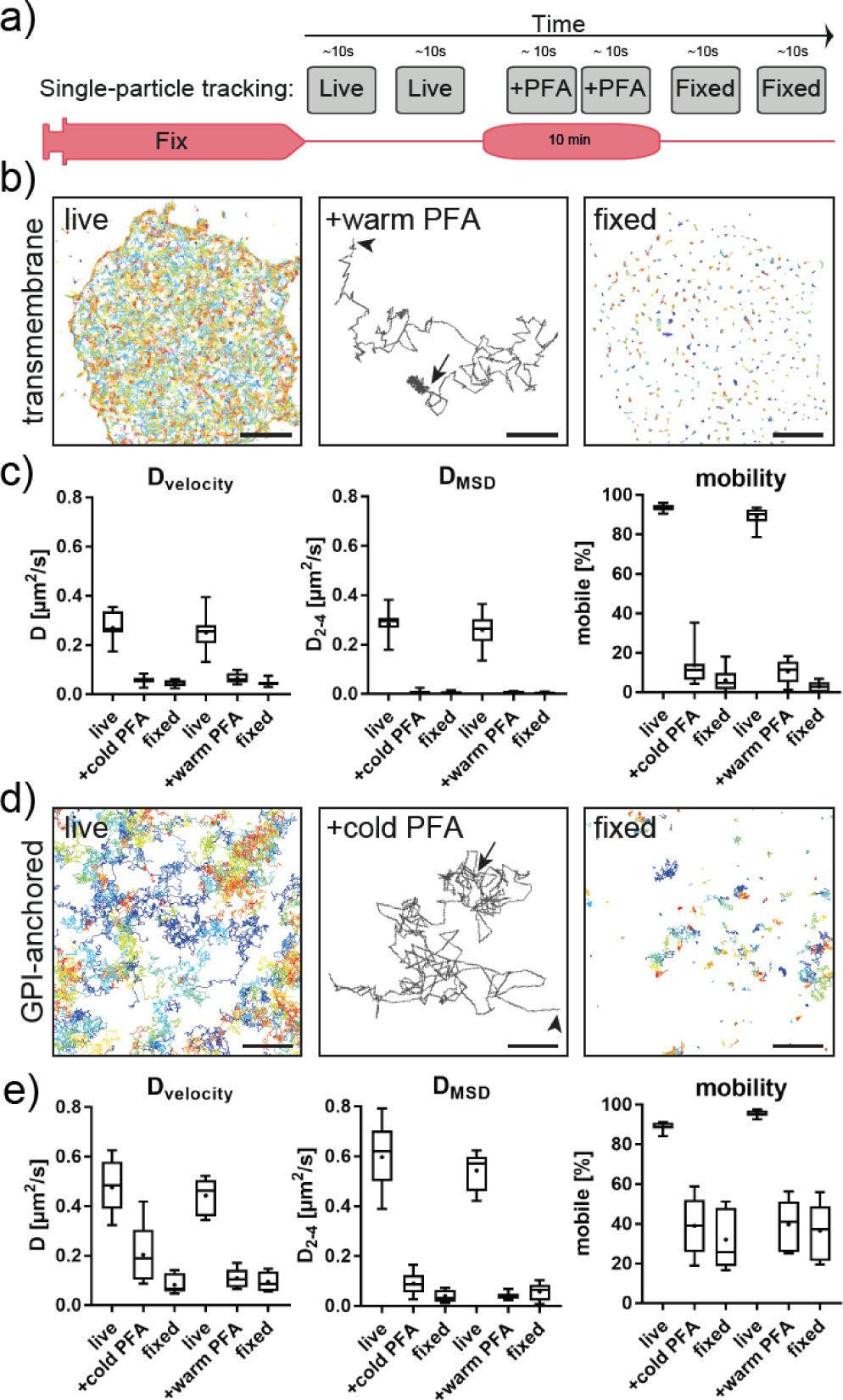
Single-particle tracking of membrane probes during live fixation. a) Experimental workflow for live and fixed cell single-particle tracking. b) Trajectories of a transmembrane probe with cytosolic and extracellular domains, tracked via fluorescently-labelled nanobodies on live cells (left). Middle panel show a typical trajectory with Brownian motion at the start (arrowhead) and immobilization upon addition of 4% PFA in PEM buffer (arrow). All transmembrane proteins appear immobilized in fixed cells. c) Quantification of diffusion coefficient D based on velocity (left), mean-square displacement (middle) and percentage of mobile (D>0.05 µm^2^/s) particles (right). No significant difference between chemical fixation at 4°C or 37°C was observed. d) Trajectories of GPI-anchored probe tracked via fluorescently labelled nanobodies on live cells (left). Middle panel show a typical trajectory with Brownian motion at the start (arrowhead) and reduced mobility upon addition of 4% PFA in PEM buffer (arrow). Some GPI-anchored proteins are immobilized in fixed cells while a fraction remains mobile. e) Quantification of diffusion coefficient D based on velocity (left), mean-square displacement (middle) and percentage of mobile (D>0.05 µm^2^/s) particles (right). No significant difference between chemical fixation at 4°C or 37°C was observed despite a trend towards faster fixation at warmer temperatures. Scale bars are 5 µm (left, right) and 500 nm (middle panels).

Thus, changes in diffusive behaviour were more dependent on the type of membrane protein tracked, rather than the fixation conditions (Fig. 4c,e), which is in agreement with previous publications (25). However, even with only 4% PFA and without any cross-linking fixatives, we observed a rapid immobilization of transmembrane proteins that would prevent artificial clustering by subsequent antibody labeling approaches.

## Discussion

The super-resolution revolution in optical microscopy offers even inexperienced users up to 10-fold increased resolution on commercial systems that have become commonly available through imaging facilities. However, established sample preparation protocols that were previously acceptable may be inadequate for super-resolution microscopy, as the inaccuracies are no longer masked by the diffraction limit. While the importance of careful sample preparation is readily accepted, its assessment remains challenging. Neglecting to recognize this cost associated with increased resolution could render imaging results useless or worse might incorrectly inform researchers about a biological system. To demonstrate sample preparation inadequacies in imaging regimes, we took advantage of NanoJ-Fluidics (28) and NanoJ-SQUIRREL (21) to compare the pre- and post-fixation actin structures and CD4 cellular organization, in the same cells. We asked what would be the influence of chemical fixation using different imaging regimes with increasing resolution (TIRF, SIM and SMLM) by correlating pre- and post-fixation images. The actin cytoskeleton acts as a scaffold that supports the plasma membrane and is in many ways involved in the organization of the plasma membrane (32, 33). However, while actin filaments are strongly affected by chemical fixation conditions, the plasma membrane itself is affected to a lesser extent. Chemical fixation is usually fast and even a simple protocol can achieve structural preservation of the organization of transmembrane proteins in the plasma membrane. Despite the availability of chemical fixation protocols that preserve the actin cytoskeleton, the predominant approach for studying protein organization is fixation with 4% PFA in PBS. Our data suggests this is insufficient to produce reliable imaging data on receptor distributions for imaging modalities that break the diffraction limit. The chemical fixation protocol used was shown to play a crucial role on the introduction of artefacts. We applied SQUIRREL, a recently developed quality metric tool (21), to quantify how much cytoskeletal structures are distorted by chemical fixation at exemplary conditions. Our approach is widely applicable to determine the impact of any fixation protocol beyond those tested. Of course, a correlation between pre- and postfixation structures is required which, albeit greatly facilitated by NanoJ-Fluidics (28), is still a time-consuming quality control approach. However, in our opinion, the benefit of increased confidence in light microscopy data is worth the added effort. In particular in challenging fields, such as plasma membrane organization where the spatial distribution of immunomodulatory receptors is directly linked to states of activity, require careful controls. The increase in cluster size and density we observed could be due to: 1) disruption of the actin cytoskeleton organization that could affect to CD4 membrane organization via protein-protein interaction; 2) fixation-induced changes in membrane properties, which would cause artificial reorganization of membrane proteins; 3) a combination of both factors. Using super-resolution microscopy we could show that the changes in CD4 organization coincided with a disrupted actin cytoskeleton profile. The cluster size in optimal conditions suggests CD4 may be organized in dimers (as seen by the mean cluster size of~ 60 nm), which is consistent with its suggested capacity to homo-dimerize, a process that may increase the avidity of its binding to MHCII (34). Further, the cluster density suggests a homogeneous distribution consistent with COS7 non-native expression. It is important to highlight that these considerable differences are in a system where CD4 does not normally exist, hence lacking the regulatory machinery or native interactions that may normally regulate CD4 distribution. Presumably, the observed differences would be more striking in CD4-positive immune cells where CD4 is linked to p56/LCK (35). Interestingly, the degree of actin cytoskeleton disruption is consistent with the extent of the changes we observe in CD4 membrane organization. After chemical fixation at 4°C we observed almost complete disruption whereas at 23°C the cell displays a mixture of regions with disrupted and non-disrupted actin structures. This suggests that despite CD4 not existing in COS7 cells in native conditions, CD4 organization is affected by the structure of the dense actin cortex (possibly through its cytoplasmic domain). Consequently, inadequate actin chemical fixation regimes can affect CD4 membrane organization and influence the biological information extracted from SMLM CD4 analysis. This is further supported by single-particle tracking experiments.

Single-particle tracking of transmembrane proteins and a GPI-anchored protein showed that the size and orientation in the plasma membrane was more important than fixation conditions. GPI-anchored proteins that reside in the outer leaflet of the plasma membrane with only indirect interaction with the submembrane cytoskeleton (36) remain largely mobile in ideal actin-preserving conditions. Any distribution or clustering analysis must rule out post-fixation aggregation, e.g. by the use of single-binders such as nanobodies. In contrast, transmembrane proteins with a cytosolic domain such as CD4 or our artificial transmembrane probe are quickly immobilized, indicating an interaction with the submembrane cytoskeleton. We also observed a trend towards faster fixation at elevated temperature. Our observation that CD4 membrane organization is affected by poor actin chemical fixation should serve as a cautionary tale for sample preparation approaches to study membrane proteins. Optimal fixation approaches preserve the cortical actin cytoskeleton structure and the organization of transmembrane proteins in a near-native state (Fig. 5). Conversely, suboptimal fixation conditions induce deformations of membrane and cytoskeleton that can result in artefacts that can influence the organization of membrane proteins, such as CD4 (Fig. 5). Although, we and others (37–40) suggest that the actin cytoskeleton and protein-protein interactions are important for membrane proteins organization, many studies using SMLM focus on imaging unknown structures and distributions of proteins that do not have a known organization. Hence, when performing essential protocol optimization, preservation of the overall cellular structure should be a priority. This work also aims to highlight that there are already established protocols that serve as excellent starting points (22, 23, 27), hardware that permits the optimization of such protocols to be streamlined (28, 41) and tools that permit seamless analysis of possible bottlenecks (21, 41). In conclusion, to extract the most from SMLM experiments it is essential to use reliable and repeatable imaging protocols that preserve, as much as possible, the overall cellular structure.

**Fig. 5.**
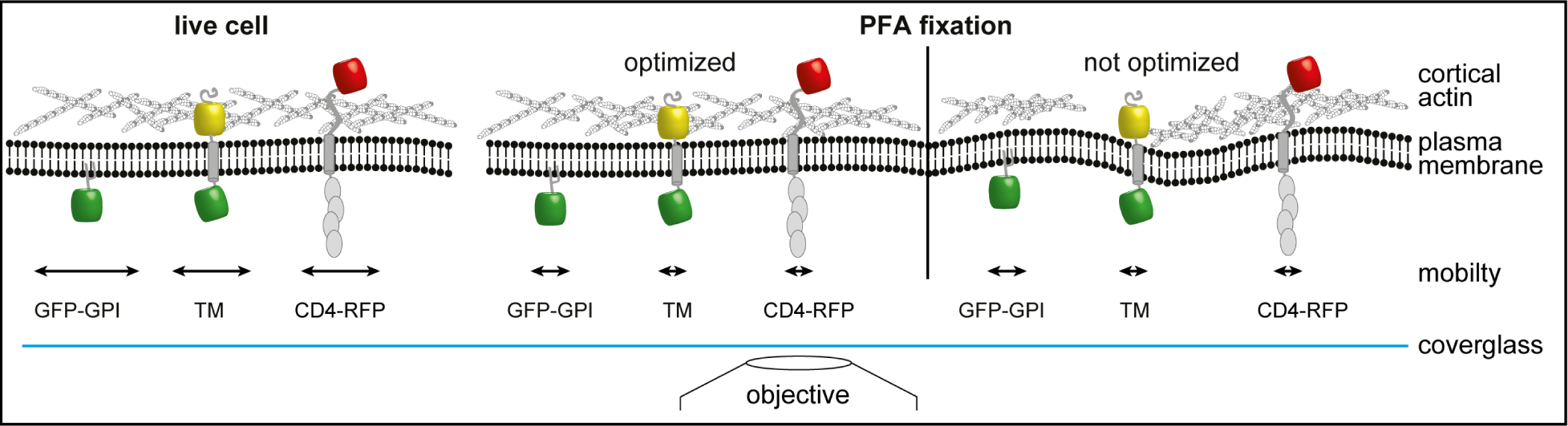
Model of changes induced by chemical fixation on membrane architecture. Optimized fixation with PFA preserves the cortical actin cytoskeleton structure in a state resembling live imaging and rapidly stops diffusion of transmembrane proteins. Suboptimal fixation conditions induce deformations of membrane and cytoskeleton and could thereby introduce artefacts. While the mobility of membrane probes is reduced similarly to optimized chemical fixation the overall organization could be altered due to interruptions of the cytoskeleton. GPI anchored GFP (GFP-GPI); artificial transmembrane protein with cytosolic and extracellular domains (mHoneydew and YFP, respectively - TM); CD4 fused to TagRFP-T (CD4-RFP).

## DATA AVAILABILITY

All data and materials used in the analysis will be made available to any researcher for purposes of reproducing or extending the analysis upon reasonable request.

## ACKNOWLEDGEMENTS

We thank Dr. Christophe Leterrier at the Neuropathophysiology Insitute (INP, CNRS-Aix Marseille University UMR 7051) for critical reading and advice. We thank the MRC-LMCB light microscopy facility for the equipment maintenance and users training. This work was funded by grants from the UK Biotechnology and Biological Sciences Research Council (BB/M022374/1; BB/P027431/1; BB/R000697/1; BB/S507532/1) (R.H., P.M.P. and C.J.); the UK Medical Research Council (MR/K015826/1) (R.H., J.M and M.M.); the Wellcome Trust (203276/Z/16/Z) (R.H); the MRC Programme Grant (MC_UU12018/7) (J.M.); the European Research Council (649101–UbiProPox) (J.M.); the MRC Programme Grant (MC_U12016/1) (M.M.); D.A. is presently a Marie Curie fellow (Marie Sklodowska-Curie grant agreement No 750673); C.J. was funded by a Commonwealth scholarship, funded by the UK government.

## AUTHOR CONTRIBUTIONS

These contributions follow the Contributor Roles Taxonomy guidelines:. Conceptualization: P.M.P., D.A., C.J., R.H.; Data curation: P.M.P., D.A., C.J.; Formal analysis: P.M.P., D.A., C.J.; Investigation: P.M.P., D.A., C.J.; Methodology: P.M.P., D.A., C.J.; Resources: P.M.P., D.A., C.J., J.M., M.M., R.H.; Validation: P.M.P., D.A., C.J., J.M., M.M., R.H.; Visualization: P.M.P., D.A., C.J.; Writing original draft: P.M.P., D.A., C.J.; Writing, review & editing: P.M.P., D.A., C.J., J.M., M.M., R.H.; Funding acquisition: P.M.P., D.A., J.M., M.M., R.H.; Project administration: P.M.P., D.A., R.H.; Supervision: P.M.P., D.A., J.M., M.M., R.H.

## Methods

### Cell lines

COS7 cells were cultured in phenol-red free DMEM (Gibco) supplemented with 2 mM GlutaMAX (Gibco), 50 U/ml penicillin, 50 µg/ml streptomycin (Penstrep, Gibco) and 10% fetal bovine serum (FBS; Gibco). Cells were grown at 37°C in a 5% CO_2_ humidified incubator. Cell lines have not been authenticated.

### Plasmids

The plasmid expressing the calponin homology domain of utrophin fused to GFP (GFP-UtrCH) was a gift from William Bement (1) (Addgene plasmid #26737). The plasmid expressing the cluster of differentiation 4 (CD4) fused to TagRFP-T25 was constructed for this study and it is available from Addgene (Addgene plasmid #119238). The plasmid expressing GPI-GFP was a kind gift from Ari Helenius. The plasmid expressing the artificial transmembrane probe was constructed based on Patrick Keller’s L-YFPGT46 (2) by adding the beta-barrel fluorophore mHoneydew on the cytosolic side to increase size (3).

### Live-to-Fixed Super-Resolution imaging

The NanoJFluidics syringe pump array was installed on a Zeiss Elyra PS.1 microscope equipped with 405, 488, 561 and 642 nm lasers (50, 200, 200 and 160 mW at the optical fibre output). All steps after cell transfection were performed on the microscope, using NanoJ-Fluidics (4, 5). COS7 cells (kind gift from Dr. A. Saiardi) were seeded on ultraclean (6) 25 mm diameter thickness 1.5H coverslips (Marienfeld) at a density of 0.3–0.9×105 cells/cm2. One day after splitting, cells were transfected with UtrCH-GFP and pCD4-TagRFP-T using Lipofectamine 2000 (Thermo Fisher Scientific) according to the manufacturer’s recommendations. Cells were imaged 1-2 days post transfection in culture medium using an Attofluor cell chamber (ThermoFisher), covered with the lid of a 35 mm dish (ThermoFisher), that was kept in place using black non-reflective aluminium tape (T205-1.0 AT205, THORLABs).

Cells were fixed at 4°C, 23°C or 37°C for 15 minutes with freshly prepared 4% paraformaldehyde (PFA) in the cytoskeleton-preserving buffer “PIPES-EGTA-Magnesium” (PEM: 80 mM PIPES pH 6.8, 5 mM EGTA, 2 mM MgCl2) (7) or at 37°C for 15 minutes with 4% PFA in Phosphate Buffer Saline (PBS: 0.14 M NaCl, 10 mM NaH2PO4, 10 mM Na2HPO4).

For stained cells (Fig. 2), after fixation cells were permeabilised (PEM with 0.25% Triton-X-100) for 20 min (at 23°C), blocked with blocking buffer (5% Bovine Serum Albumin (BSA) in PEM) for 30 minutes (at 23°C), and stained with anti-CD4 mAb (OKT4, 6 µg/ml) for 60 min (at 23°C), followed by anti-mouse Alexa Fluor 568 secondary Ab (Molecular Probes) for 60 min (at 23°C). Structured Illumination Microscopy (SIM) imaging was performed using Plan-Apochromat 63x/1.4 oil DIC M27 objective, in a Zeiss Elyra PS.1 microscope (Zeiss). Images were acquired using 5 phase shifts and 3 grid rotations with the 561 nm and 488 nm lasers (at 5-10%of maximum output), and filter set 4 (1851-248, Zeiss). Images were acquired using a sCMOS (pco.edge sCMOS) camera. Total Internal Reflection Fluorescence (TIRF) SRRF imaging of live COS7 cells was performed at 37 °C and 5% CO2 on a Zeiss Elyra PS.1 microscope, by acquiring 100 frames (33 FPS) with 488 nm and 561 nm laser illumination at 0.5% of maximum output. A 100x TIRF objective (Plan-APOCHROMAT 100x/1.46 Oil, Zeiss) with additional 1.6x magnification was used to collect fluorescence onto an EMCCD camera (iXon Ultra 897, Andor), yielding a pixel size of 100 nm. TIRF STORM imaging of anti-CD4 Alexa Fluor 568 in fixed cells was performed on the same system. 50,000 frames were acquired with 33 ms exposure and 561 nm laser illumination at maximum output power with 405 nm pumping when required (0.5-1% of maximum output when the blinking density was bellow 1 particle/µm2). STORM imaging was performed in GLOX buffer (150mM Tris, pH 8, 1% glycerol, 1% glucose, 10mM NaCl, 1% β-mercaptoethanol, 0.5 mg/ml glucose oxidase, 40 µg/ml catalase). Single-particle tracking was performed in medium at 37°C and 5% CO2 on a Zeiss Elyra PS.1 microscope in TIRF mode by acquiring 250/500 frames at 45 FPS with 642 nm laser illumination at 5% of maximum output. For live-fixation, medium was replaced by either ice-cold or prewarmed (at 37°C) 4% PFA in PEM buffer.

### Image reconstruction and analysis

For Fig. 2 images were processed using the ZEN software (2012, version 8.1.6.484, Zeiss). For channel alignment, a multi-coloured bead slide was imaged using the same image acquisition settings. For STORM datasets localizations were detected and rendered using ThunderSTORM (8) with default settings. Fourier Ring Correlation (FRC) values were obtained using NanoJ-SQUIRREL after reconstruction of original data separated into two different stacks composed of odd or even images (9). NanoJ-SQUIRREL and ThunderSTORM are available in Fiji (10). Statistical analysis (ordinary one-way ANOVA) was performed using Prism7 (GraphPad). Single-particle tracking data was analysed using Trackmate (11) in Fiji and MSDanalyzer (12) in MATLAB (Mathworks).

## Bibliography

1 Eric Betzig, George H Patterson, Rachid Sougrat, O Wolf Lindwasser, Scott Olenych, Juan S Bonifacino, Michael W Davidson, Jennifer Lippincott-Schwartz, and Harald F Hess. Imaging intracellular fluorescent proteins at nanometer resolution. Science, 313(5793): 1642–1645, 2006.

2 Michael J Rust, Mark Bates, and Xiaowei Zhuang. Sub-diffraction-limit imaging by stochastic optical reconstruction microscopy (storm). Nature methods, 3(10): 793, 2006.

3 Philip R Nicovich, Dylan M Owen, and Katharina Gaus. Turning single-molecule localization microscopy into a quantitative bioanalytical tool. Nature protocols, 12(3): 453, 2017.

4 Florian Levet, Eric Hosy, Adel Kechkar, Corey Butler, Anne Beghin, Daniel Choquet, and Jean-Baptiste Sibarita. Sr-tesseler: a method to segment and quantify localization-based super-resolution microscopy data. Nature methods, 12(11): 1065, 2015.

5 Robert Gray, David Albrecht, Corina Beerli, Gary Cohen, Ricardo Henriques, and Jason Mercer. Nanoscale polarization of the vaccinia virus entry fusion complex drives efficient fusion. bioRxiv, page 360073, 2018.

6 Marta Fernández-Suárez and Alice Y Ting. Fluorescent probes for super-resolution imaging in living cells. Nature reviews Molecular cell biology, 9(12): 929, 2008.

7 Filipa B Lopes, Ŝtefan Bálint, Salvatore Valvo, James H Felce, Edith M Hessel, Michael L Dustin, and Daniel M Davis. Membrane nanoclusters of fcγri segregate from inhibitory sirpα upon activation of human macrophages. J Cell Biol, pages jcb–201608094, 2017.

8 Sinem K Saka, Alf Honigmann, Christian Eggeling, Stefan W Hell, Thorsten Lang, and Silvio O Rizzoli. Multi-protein assemblies underlie the mesoscale organization of the plasma membrane. Nature communications, 5:4509, 2014.

9 Jinmin Lee, Prabuddha Sengupta, Joseph Brzostowski, Jennifer Lippincott-Schwartz, and Susan K Pierce. The nanoscale spatial organization of b-cell receptors on immunoglobulin m–and g–expressing human b-cells. Molecular biology of the cell, 28(4): 511–523, 2017.

10 Helena Soares, Ricardo Henriques, Martin Sachse, Leandro Ventimiglia, Miguel A Alonso, Christophe Zimmer, Maria-Isabel Thoulouze, and Andrés Alcover. Regulated vesicle fusion generates signaling nanoterritories that control t cell activation at the immunological synapse. Journal of Experimental Medicine, pages jem–20130150, 2013.

11 Joana G Silva, Nuno P Martins, Ricardo Henriques, and Helena Soares. Hiv-1 nef impairs the formation of calcium membrane territories controlling the signaling nanoarchitecture at the immunological synapse. The Journal of Immunology, page 1601132, 2016.

12 Paul E Love and Sandra M Hayes. Itam-mediated signaling by the t-cell antigen receptor. Cold Spring Harbor perspectives in biology, page a002485, 2010.

13 Maro Iliopoulou, Rory Nolan, Luis Alvarez, Yasunori Watanabe, Charles A Coomer, G Maria Jakobsdottir, Thomas A Bowden, and Sergi Padilla-Parra. A dynamic three-step mechanism drives the hiv-1 pre-fusion reaction. Nature structural & molecular biology, 25(9): 814, 2018.

14 Pieta K Mattila, Facundo D Batista, and Bebhinn Treanor. Dynamics of the actin cytoskeleton mediates receptor cross talk: An emerging concept in tuning receptor signaling. J Cell Biol, 212(3): 267–280, 2016.

15 Mary L Kraft. Plasma membrane organization and function: moving past lipid rafts. Molecular biology of the cell, 24(18): 2765–2768, 2013.

16 William A Comrie and Janis K Burkhardt. Action and traction: cytoskeletal control of receptor triggering at the immunological synapse. Frontiers in immunology, 7:68, 2016.

17 Olivia L Mooren, Brian J Galletta, and John A Cooper. Roles for actin assembly in endocytosis. Annual review of biochemistry, 81, 2012.

18 William S Trimble and Sergio Grinstein. Barriers to the free diffusion of proteins and lipids in the plasma membrane. J Cell Biol, 208(3): 259–271, 2015.

19 Steffen J Sahl, Stefan W Hell, and Stefan Jakobs. Fluorescence nanoscopy in cell biology. Nature reviews Molecular cell biology, 18(11): 685, 2017.

20 Theresia EB Stradal and Mario Schelhaas. Actin dynamics in host-pathogen interaction. FEBS Letters.

21 Siân Culley, David Albrecht, Caron Jacobs, Pedro Matos Pereira, Christophe Leterrier, Jason Mercer, and Ricardo Henriques. Quantitative mapping and minimization of super-resolution optical imaging artifacts. Nature methods, 15(4): 263, 2018.

22 Daniela Leyton-Puig, Katarzyna M Kedziora, Tadamoto Isogai, Bram van den Broek, Kees Jalink, and Metello Innocenti. Pfa fixation enables artifact-free super-resolution imaging of the actin cytoskeleton and associated proteins. Biology open, 5(7): 1001–1009, 2016.

23 Donna R Whelan and Toby DM Bell. Image artifacts in single molecule localization microscopy: why optimization of sample preparation protocols matters. Scientific reports, 5: 7924, 2015.

24 Ying Zhang, Tao Huang, Danielle M Jorgens, Andrew Nickerson, Li-Jung Lin, Joshua Pelz, Joe W Gray, Claudia S López, and Xiaolin Nan. Quantitating morphological changes in biological samples during scanning electron microscopy sample preparation with correlative super-resolution microscopy. PLoS One, 12(5):e0176839, 2017.

25 Kenji AK Tanaka, Kenichi GN Suzuki, Yuki M Shirai, Shusaku T Shibutani, Manami SH Miyahara, Hisae Tsuboi, Miyako Yahara, Akihiko Yoshimura, Satyajit Mayor, Takahiro K Fujiwara, et al. Membrane molecules mobile even after chemical fixation. Nature Methods, 7(11): 865, 2010.

26 Ke Xu, Guisheng Zhong, and Xiaowei Zhuang. Actin, spectrin, and associated proteins form a periodic cytoskeletal structure in axons. Science, 339(6118): 452–456, 2013.

27 Katharina N Richter, Natalia H Revelo, Katharina J Seitz, Martin S Helm, Deblina Sarkar, Rebecca S Saleeb, Elisa D’Este, Jessica Eberle, Eva Wagner, Christian Vogl, et al. Glyoxal as an alternative fixative to formaldehyde in immunostaining and super-resolution microscopy. The EMBO journal, 37(1): 139–159, 2018.

28 Pedro Almada, Pedro Pereira, Siân Culley, Ghislaine Caillol, Fanny Boroni-Rueda, Christina L Dix, Romain F Laine, Guillaume Charras, Buzz Baum, Christophe Leterrier, et al. Automating multimodal microscopy with nanoj-fluidics. bioRxiv, page 320416, 2018.

29 Mats GL Gustafsson. Surpassing the lateral resolution limit by a factor of two using structured illumination microscopy. Journal of microscopy, 198(2): 82–87, 2000.

30 Brian M Burkel, George Von Dassow, and William M Bement. Versatile fluorescent probes for actin filaments based on the actin-binding domain of utrophin. Cell motility and the cytoskeleton, 64(11): 822–832, 2007.

31 Gregory Giannone, Eric Hosy, Florian Levet, Audrey Constals, Katrin Schulze, Alexander I Sobolevsky, Michael P Rosconi, Eric Gouaux, Robert Tampé, Daniel Choquet, et al. Dynamic superresolution imaging of endogenous proteins on living cells at ultra-high density. Biophysical journal, 99(4): 1303–1310, 2010.

32 Senthil Arumugam, Eugene P Petrov, and Petra Schwille. Cytoskeletal pinning controls phase separation in multicomponent lipid membranes. Biophysical journal, 108(5): 1104–1113, 2015.

33 Sven Kenjiro Vogel, Ferdinand Greiss, Alena Khmelinskaia, and Petra Schwille. Control of lipid domain organization by a biomimetic contractile actomyosin cortex. Elife, 6:e24350, 2017.

34 Hao Wu, Peter D Kwong, and Wayne A Hendrickson. Dimeric association and segmental variability in the structure of human cd4. Nature, 387(6632): 527, 1997.

35 Annegret Pelchen-Matthews, Isabelle Boulet, Dan R Littman, Remi Fagard, and Mark Marsh. The protein tyrosine kinase p56lck inhibits cd4 endocytosis by preventing entry of cd4 into coated pits. The Journal of cell biology, 117(2): 279–290, 1992.

36 Riya Raghupathy, Anupama Ambika Anilkumar, Anirban Polley, Parvinder Pal Singh, Mahipal Yadav, Charles Johnson, Sharad Suryawanshi, Varma Saikam, Sanghapal D Sawant, Aniruddha Panda, et al. Transbilayer lipid interactions mediate nanoclustering of lipid-anchored proteins. Cell, 161(3): 581–594, 2015.

37 Manasa V Gudheti, Nikki M Curthoys, Travis J Gould, Dahan Kim, Mudalige S Gunewardene, Kristin A Gabor, Julie A Gosse, Carol H Kim, Joshua Zimmerberg, and Samuel T Hess. Actin mediates the nanoscale membrane organization of the clustered membrane protein influenza hemagglutinin. Biophysical journal, 104(10): 2182–2192, 2013.

38 Herlinde De Keersmaecker, Rafael Camacho, David Manuel Rantasa, Eduard Fron, Hiroshi Uji-i, Hideaki Mizuno, and Susana Rocha. Mapping transient protein interactions at the nanoscale in living mammalian cells. ACS nano, 2018.

39 Sanaz Sadegh, Jenny L Higgins, Patrick C Mannion, Michael M Tamkun, and Diego Krapf. Plasma membrane is compartmentalized by a self-similar cortical actin meshwork. Physical Review X, 7(1): 011031, 2017.

40 George W Ashdown, Garth L Burn, David J Williamson, Elvis Pandžić, Ruby Peters, Michael Holden, Helge Ewers, Lin Shao, Paul W Wiseman, and Dylan M Owen. Live-cell super-resolution reveals f-actin and plasma membrane dynamics at the t cell synapse. Biophysical journal, 112(8): 1703–1713, 2017.

41 Romain Laine, Kalina Tosheva, Nils Gustafsson, Robert DM Gray, Pedro Almada, David Albrecht, Gabriel T Risa, Fredrik Hurtig, Ann-Christin Lindås, Buzz Baum, et al. Nanoj: a high-performance open-source super-resolution microscopy toolbox. bioRxiv, page 432674, 2018.

## Bibliography

1 Brian M Burkel, George Von Dassow, and William M Bement. Versatile fluorescent probes for actin filaments based on the actin-binding domain of utrophin. Cell motility and the cytoskeleton, 64(11): 822–832, 2007.

2 Patrick Keller, Derek Toomre, Elena Díaz, Jamie White, and Kai Simons. Multicolour imaging of post-golgi sorting and trafficking in live cells. Nature cell biology, 3(2): 140, 2001.

3 David Albrecht, Christian M Winterflood, and Helge Ewers. Dual color single particle tracking via nanobodies. Methods and applications in fluorescence, 3(2): 024001, 2015.

4 Pedro Almada, Pedro Pereira, Siân Culley, Ghislaine Caillol, Fanny Boroni-Rueda, Christina L Dix, Romain F Laine, Guillaume Charras, Buzz Baum, Christophe Leterrier, et al. Automating multimodal microscopy with nanoj-fluidics. bioRxiv, page 320416, 2018.

5 Christina L Dix, Helen K Matthews, Marina Uroz, Susannah McLaren, Lucie Wolf, Nicholas Heatley, Zaw Win, Pedro Almada, Ricardo Henriques, Michael Boutros, et al. The role of mitotic cell-substrate adhesion re-modeling in animal cell division. Developmental cell, 45 (1): 132–145, 2018.

6 Pedro M Pereira, Pedro Almada, and Ricardo Henriques. High-content 3d multicolor super-resolution localization microscopy. In Methods in cell biology, volume 125, pages 95–117. Academic Press, 2015.

7 Daniela Leyton-Puig, Katarzyna M Kedziora, Tadamoto Isogai, Bram van den Broek, Kees Jalink, and Metello Innocenti. Pfa fixation enables artifact-free super-resolution imaging of the actin cytoskeleton and associated proteins. Biology open, 5(7): 1001–1009, 2016.

8 Martin Ovesnỳ, Pavel Křížek, Josef Borkovec, Zdeněk Ŝvindrych, and Guy M Hagen. Thunderstorm: a comprehensive imagej plug-in for palm and storm data analysis and super-resolution imaging. Bioinformatics, 30(16): 2389–2390, 2014.

9 Siân Culley, David Albrecht, Caron Jacobs, Pedro Matos Pereira, Christophe Leterrier, Jason Mercer, and Ricardo Henriques. Quantitative mapping and minimization of super-resolution optical imaging artifacts. Nature methods, 15(4): 263, 2018.

10 Johannes Schindelin, Ignacio Arganda-Carreras, Erwin Frise, Verena Kaynig, Mark Longair, Tobias Pietzsch, Stephan Preibisch, Curtis Rueden, Stephan Saalfeld, Benjamin Schmid, et al. Fiji: an open-source platform for biological-image analysis. Nature methods, 9(7): 676, 2012.

11 Jean-Yves Tinevez, Nick Perry, Johannes Schindelin, Genevieve M Hoopes, Gregory D Reynolds, Emmanuel Laplantine, Sebastian Y Bednarek, Spencer L Shorte, and Kevin W Eliceiri. Trackmate: An open and extensible platform for single-particle tracking. Methods, 115:80–90, 2017.

12 Nadine Tarantino, Jean-Yves Tinevez, Elizabeth Faris Crowell, Bertrand Boisson, Ricardo Henriques, Musa Mhlanga, Fabrice Agou, Alain Israël, and Emmanuel Laplantine. Tnf and il-1 exhibit distinct ubiquitin requirements for inducing nemo–ikk supramolecular structures. J Cell Biol, 204(2): 231–245, 2014.

